# CD8+ T cells and Humoral Immunity Influence the Development of Antibody-Dependent Enhancement: Implications for Vaccine Design

**DOI:** 10.64898/2026.03.10.710878

**Authors:** Riley Drake, Hasan Ahmed, Anmol Chandele, Rafi Ahmed, Kaja Murali-Krishna, Rustom Antia

## Abstract

Dengue virus (DENV) infection is a major and growing global health threat. The development of a safe and effective vaccine regardless of prior dengue immunity remains an unmet need. Virus-specific antibodies typically limit viral replication. In contrast, low-to-intermediate dengue-specific antibody titers can enhance viral replication and increase disease severity—a phenomenon known as antibody-dependent enhancement (ADE). ADE complicates dengue vaccine development; signals of ADE were observed in the CYD-TDV (Dengvaxia) vaccine trials but have not yet been reported for trials of TAK-003 (Qdenga). To better understand how prior immunity influences dengue infection outcomes, we developed a mechanistic within-host model of acute dengue infection dynamics that incorporates both humoral immunity and CD8+ T cells. Model simulations predict that severe disease is most likely when dengue-specific antibody titers are intermediate and dengue-specific CD8+ T-cell immunity is low at the time of infection. In our simulations, increasing pre-infection levels of CD8+ T-cell immunity reduces disease severity in a dose-dependent manner and can mitigate ADE. Our results suggest a mechanistic interaction between antibody levels and CD8+ T-cell immunity that may influence whether enhanced viral replication leads to severe disease. The model provides a possible explanation for why ADE is frequently observed in infections of nonhuman primates following passive antibody transfer, yet is less common during secondary infections in humans with memory CD8+ T cells. These findings may help interpret differences in reported clinical outcomes of dengue vaccine trials and highlight the importance of considering CD8+ T-cell responses in the design of future dengue vaccines.

**Importance:** Vaccines, including those against dengue viruses, are typically designed to elicit humoral (antibody-mediated) immunity, as antibody titers are commonly used as primary correlates of protection. Because CD8+ T-cell responses are not usually primary correlates and are more challenging to measure, they have received comparatively less attention. For viruses such as dengue, however, antibodies can exacerbate disease severity through antibody-dependent enhancement (ADE), underscoring the need to understand how humoral and cellular immunity interact. Here, we present a mechanistic model showing that severe dengue disease arises not only from low-to-intermediate antibody levels but also from insufficient CD8+ T-cell immunity at the time of infection. The model predicts that CD8+ T-cell responses can mitigate ADE-associated pathology and may help explain the differing levels of protection observed for CYD-TDV and TAK-003 vaccines. These findings suggest that evaluating CD8+ T-cell responses alongside antibody titers may be important for assessing vaccine safety and protection.

## 1 Introduction

Dengue is a disease of global importance, with over 14 million reported cases globally in 2024; about 50% of the world’s population lives in areas where dengue transmission occurs [18]. Specific antibodies against most viruses attenuate or abrogate viral replication. In contrast, a characteristic feature of dengue virus is antibody-dependent enhancement (ADE), in which low-to-intermediate titers of preexisting antibody enhance, rather than attenuate, viral replication, leading to increased disease severity [20, 21, 22, 23, 25, 26, 12]. Consequently, as shown in Fig. 1, viral replication and infection pathology may be highest at intermediate antibody titers and subsequently decrease at higher antibody concentrations.

**Figure 1.**
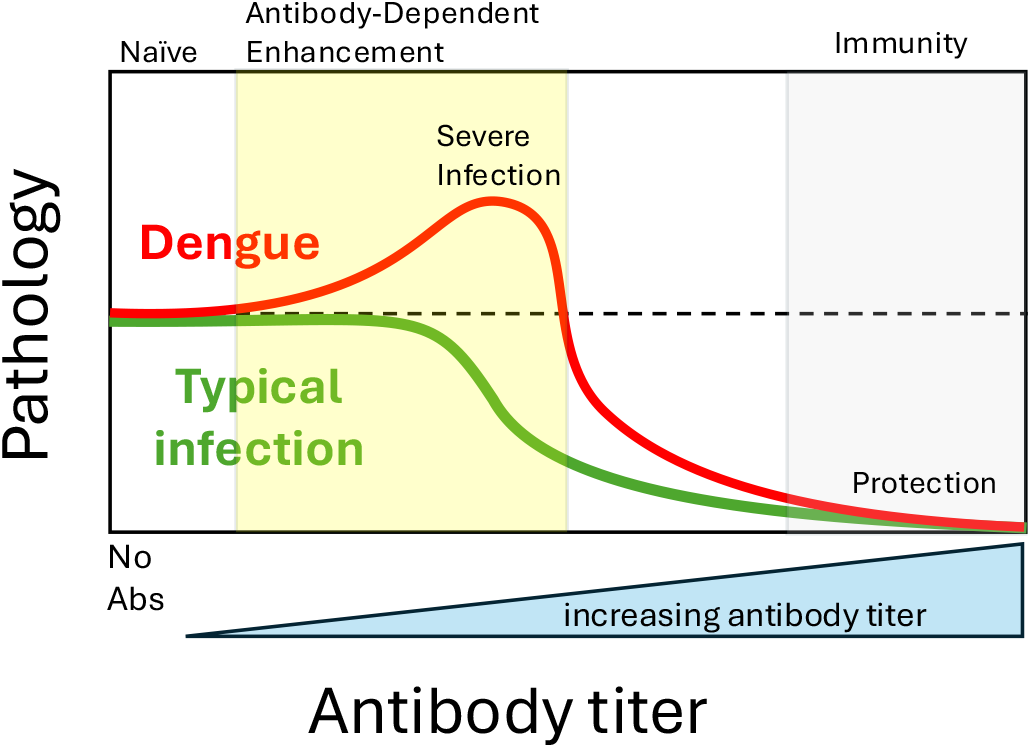
Virus-specific antibodies typically limit viral replication and limit infection pathology. In contrast, for the dengue virus, a low antibody titer can enhance virus replication and increase the severity of infection—a phenomenon known as antibody-dependent enhancement (ADE). High antibody levels clear the virus and confer protection against subsequent reinfection.

The strongest evidence for antibody-dependent enhancement of dengue virus replication comes from two types of experimental studies: cell culture experiments that measure virus replication *in vitro*, and nonhuman primate (NHP) studies that assess viral replication *in vivo* after passive antibody transfer and experimental infection. In the *in vitro* studies, the addition of intermediate amounts of antibody enhanced viral replication [16, 12]. In the NHP studies, passive transfer of intermediate amounts of anti-dengue antibodies to NHPs prior to infection resulted in a 10-to 100-fold enhancement of virus replication [16]. In both the *in vitro* and passive transfer settings, virus replication is abrogated at high antibody concentrations [16, 12]. Epidemiological studies of natural dengue infection have identified associations consistent with ADE, showing that individuals with low or intermediate titers of dengue-specific antibodies are at higher risk of severe disease than antibody-naive individuals or those with high antibody titers [29, 43]. Of note, unlike in the NHP studies described earlier where enhancing titers of antibodies resulted in a robust increase in virus replication, in the human studies only a small fraction of individuals with antibody titers in the enhancing range developed severe disease [29, 43].

ADE has important implications for vaccination. The first two licensed dengue vaccines, CYD-TDV and TAK-003, were designed to elicit robust antibody responses against all four dengue virus serotypes [27]. Both vaccines use live-attenuated recombinant vectors to express the surface proteins of the four dengue serotypes. CYD-TDV and TAK-003 differ in the non-structural proteins that form the backbone of the live attenuated virus. The backbone of CYD-TDV (Dengvaxia) is from the yellow fever virus (YFV-17D); in contrast, the backbone of the TAK-003 (Qdenga) vaccine is derived from an attenuated DENV-2 strain (16681-PDK53) [10]. Both vaccines elicit robust antibody responses that are initially protective [44]. However, as antibody titers waned, a small increase in the risk of severe disease was observed among children immunized with Dengvaxia [3, 56], with a more pronounced effect in seronegative recipients [45]. Following these findings, the use of Dengvaxia in seronegative individuals was discontinued. TAK-003, which elicited robust long-lived CD8+ T-cell responses against DENV-2 [54, 32], did not increase the risk of severe disease [37] and is in widespread use [41, 49]. The cause of the different outcomes in these two vaccine trials is not known.

To generate a more nuanced understanding of the dynamics of virus and immunity following dengue infections, we developed a mechanistic model of virus infection and immunity dynamics. Like previously published mechanistic models of ADE [33], our model explicitly includes innate and humoral immunity. Here we extend this framework by incorporating CD8+ T-cell responses, which allows us to explore how the interplay between humoral and cellular immunity determines the outcome of dengue infection and the risk of ADE. The model recapitulates *in vitro* and NHP studies described earlier, which show enhanced virus replication at intermediate antibody concentrations. The model provides a possible explanation for the different outcomes of the two dengue vaccines. It also suggests a contributing factor for why severe disease caused by ADE is a relatively rare outcome of human infection. Finally, we consider implications for the design of future dengue vaccines.

## 2 Results

We built a mechanistic model to describe the within-host dynamics of dengue infections. The model builds on earlier models that track populations of uninfected and infected cells, virus, and humoral immune responses [33] by the addition of CD8+ T-cell responses. Fig. 2 shows the model’s major components. Briefly, following infection, the growth of the viral population rapidly triggers innate immunity, and at a slower rate, humoral and CD8+ T-cell responses. As in an earlier study, sub-neutralizing antibody concentrations enhance viral replication [33]. In keeping with previous clinical observations [51, 53], we assume that the pathology of dengue infection is proportional to infectious virus load (*V*_0_ + *V*_1_), which we quantify by the area under the virus titer. Parameter values were chosen to reproduce the approximate timescale and viral load dynamics observed in experimental dengue infections and challenge studies. A more detailed description of the model and the equations used is provided in the Materials and Methods, and the simulation parameters are provided in the Supplemental Material.

**Figure 2.**
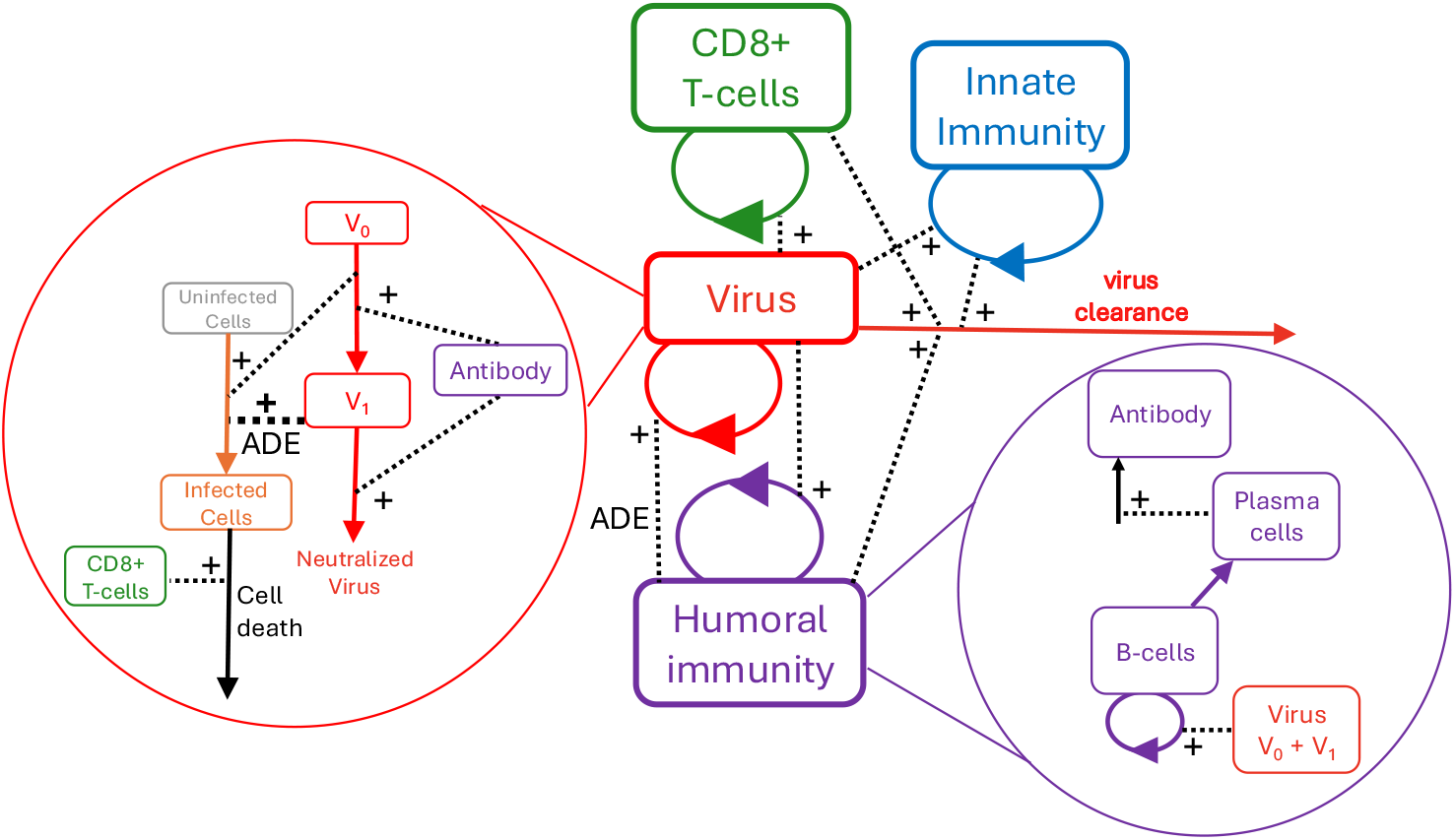
Conceptual diagram of the within-host model of dengue infection dynamics. Boxes representing virus, innate immunity, humoral immunity, and CD8+ T cells indicate the main components of the model. The effects of antibody-dependent enhancement (ADE) and neutralization are captured by tracking two viral populations: unbound virus (*V*_0_) and virus partially bound by antibody (*V*_1_). Virus in the *V*_1_ state exhibits an enhanced rate of cell infection relative to *V*_0_. When sufficient antibody binds, virus is fully neutralized, rendered unable to infect cells, and rapidly cleared. Humoral immunity includes populations of B cells, plasma cells, and secreted antibody. The proliferation of both B cells and CD8+ T cells is dependent on virus concentration. See Materials and Methods for details.

In Fig. 3 we show the results of a simulation of the dynamics of a primary dengue infection. Following its introduction, the virus grows rapidly. The virus quickly induces an innate immune response, which slows its growth. The virus also induces expansion of both humoral and CD8+ T-cell immunity. Around 1-2 weeks after infection, the levels of CD8+ T cells and antibodies are sufficient to control and then clear the virus. The simulated infection resolves on a timescale similar to that observed in experimental dengue challenge studies [55].

**Figure 3.**
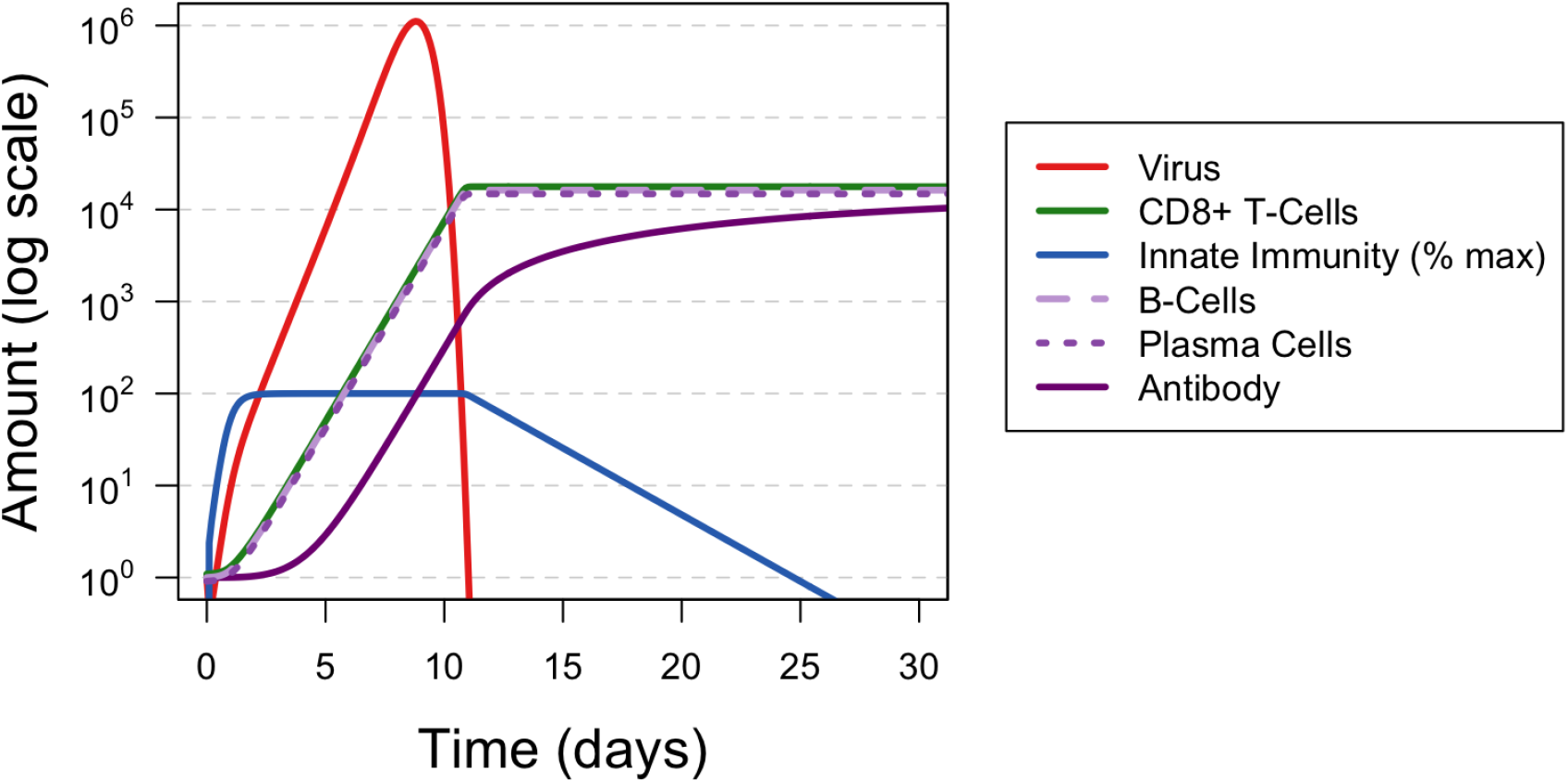
Dynamics of Primary Infection. Diagram of the dynamics of primary infection showing populations of Unbound virus (V_0_), partially bound virus (V_1_), CD8+ T cells (T), Innate Immunity (X) (X is on a different scale than other state variables and is plotted as a % of maximum level), B Cells (B), Plasma Cells (P), and antibody. Each of these state variables is shown from infection through resolution, with a time horizon of about 30 days. In the case of primary infection with no pre-existing immunity shown here, the infection (as measured by the number of infectious viral particles, *V*_0_+*V*_1_) is cleared by Day 12 post-infection. For visual clarity, overlapping immune trajectories were offset by *<*5% without altering underlying dynamics.

We then used the model to explore how infection dynamics depend on the level of prior immunity at the time of infection. Fig. 4 shows simulations across a range of initial humoral and CD8+ T-cell immunity levels, with virus (*V*_0_ + *V*_1_), CD8+ T cells, and antibody shown in red, green, and purple, respectively. The bottom-left pane of Fig. 4 reflects the dynamics of primary infection in an individual with no pre-existing immunity to dengue. The dashed black line indicates the peak amount of infectious virus reached in this scenario, which facilitates visual comparison of how prior immunity affects peak virus titer. If the infection occurs in individuals with intermediate levels of B-cell immunity but only naive levels of CD8+ T-cell immunity (plots in the middle of the bottom row), we observed higher virus replication, which characterizes ADE. We observe that higher levels of CD8+ T-cell immunity at the time of infection result in more rapid virus control, indicating that CD8+ T cells can mitigate ADE. High levels of either humoral or CD8+ T-cell immunity at the time of infection likewise promote rapid virus control. These patterns are consistent when pathology is measured by peak virus load or by the area under the curve (AUC) of the virus load, as further detailed in Fig. S7.

**Figure 4.**
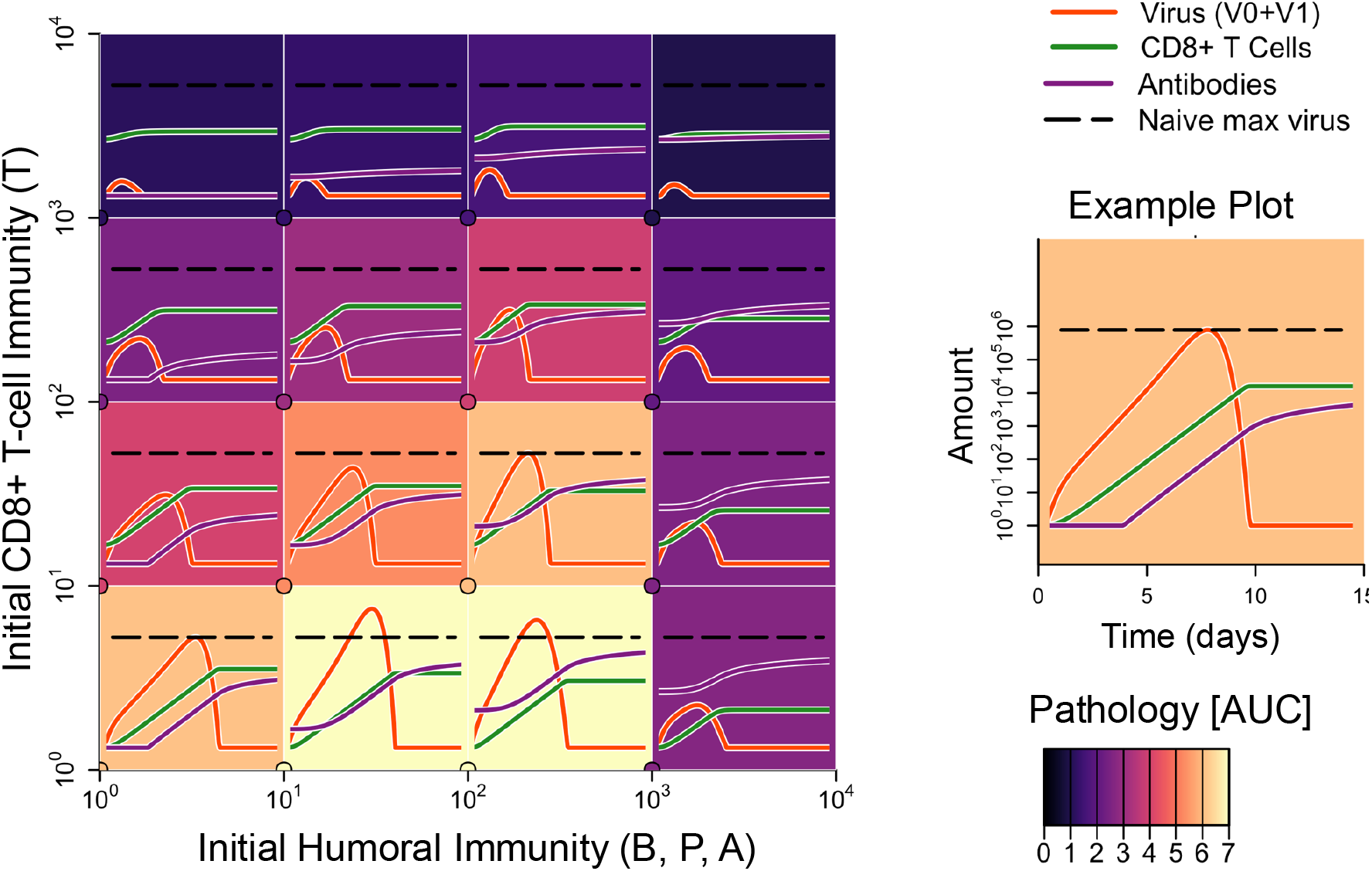
Plots showing how infection dynamics depend on the levels of humoral and CD8+ T-cell immunity at the time of infection. Each panel represents a simulated infection initiated with specific levels of humoral immunity (x-coordinate) and CD8+ T-cell immunity (y-coordinate), indicated by the circle in the lower left corner of each plot. The lines depict the dynamics of virus (red), antibody (purple), and CD8+ T-cell immunity (green) over the course of infection. The black dashed line indicates the maximum viral titer achieved during infection in a naive individual with no prior immunity. Virus titer is enhanced when infections begin with intermediate humoral immunity and naive levels of CD8+ T cells, reflecting ADE. In contrast, the presence of dengue-specific CD8+ T cells prior to infection protects against antibody-dependent enhancement in a dose-dependent manner. The background color of each panel represents the AUC of the virus load, an additional measure of pathology. Similar patterns are observed when pathology is quantified by peak virus load or by AUC, as detailed in Fig. S3.

Next, we performed a two-dimensional initial condition scan to explore how the levels of humoral and CD8+ T-cell immunity affect pathology during the secondary infection. These results are shown in Fig. 5, which displays the pathology outcome for an individual starting with different levels of humoral immunity (x-axis) and CD8+ T-cell immunity (y-axis) at the time of infection. We approximate infection severity using the area under the curve (AUC) of infectious virus over the entire course of infection. Here, we see that infections (bottom left of the heatmap) cause some pathology. In contrast, infections that start with large B-cell or CD8+ T-cell immunity prevent productive infection and cause little pathology. Finally, we observed pathology reflective of ADE at low or intermediate levels of humoral immunity, which can be mitigated by increasing CD8+ T-cell immunity.

**Figure 5.**
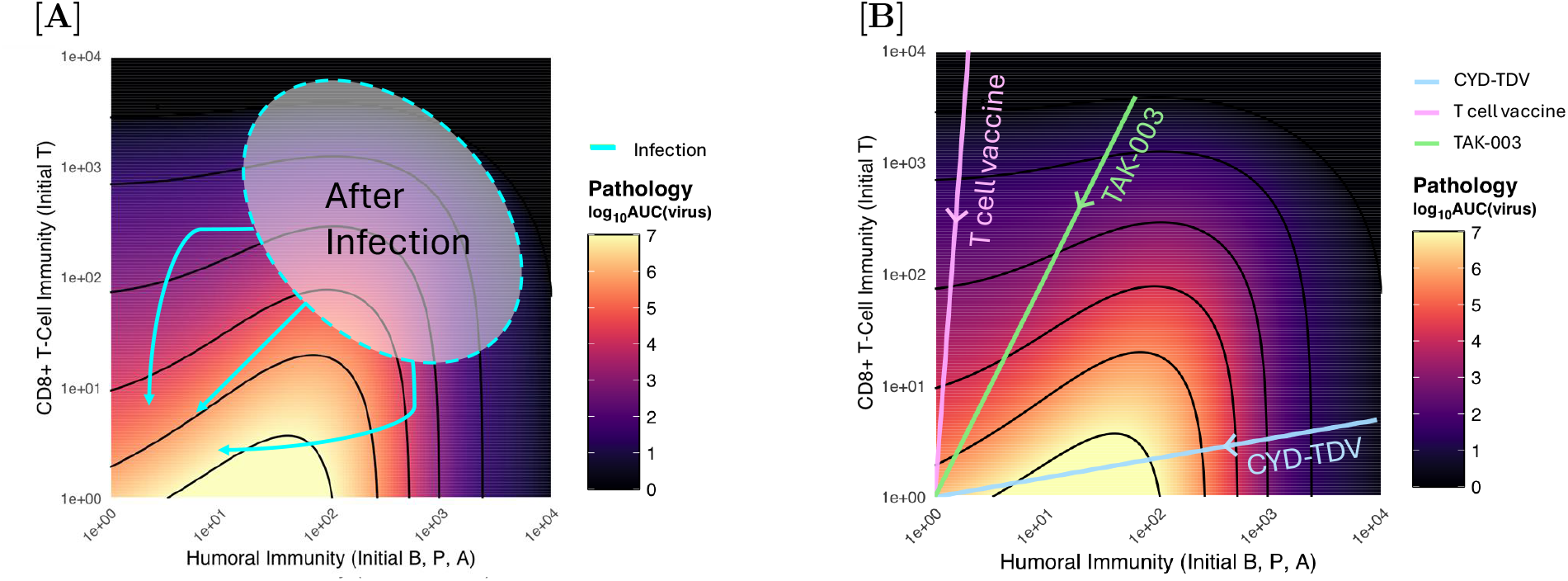
The effect of pre-existing immunity on pathology: The color on the heatmap indicates how the severity of the infection depends on the initial levels of humoral immunity (x-axis) and CD8+ T-cell immunity (y-axis). We see increased pathology when individuals begin infections with low or intermediate humoral immunity and low CD8+ T-cell immunity. Increasing the number of specific CD8+ T cells at the time of infection protects against this enhancement of pathology due to ADE in a dose-dependent manner. [A] shows possible trajectories of immune waning after infection, [B] shows possible trajectories of waning after the different vaccines. We expect Dengvaxia to elicit fewer dengue-specific CD8+ T cells because the non-structural proteins that form its backbone are from YFV, and TAK-003 to elicit higher numbers of CD8+ T cells as its backbone is from DENV-2.

### Immune Waning and Pathology of Reinfection

Infection or immunization typically elicits a robust adaptive response consisting of both humoral and CD8+ T-cell immunity. There is substantial heterogeneity among individuals in the magnitude of immune responses elicited by infection, hereafter referred to as the peak post-infection response. Protection declines following infection because both arms of the adaptive immune response will wane from the post-infection endpoint, typically rapidly in the first couple of years and more slowly thereafter [43, 42]. Substantial heterogeneity in the magnitude of humoral and CD8+ T-cell immunity following infection, as well as differences in the rates at which these responses wane, gives rise to a wide range of possible reinfection pathologies. The risk of antibody-dependent enhancement (ADE) as immunity wanes may be elevated in individuals with weak or rapidly waning CD8+ T-cell responses, whereas individuals with relatively strong CD8+ T-cell responses may experience a lower risk of severe disease across their waning trajectory (Fig. 5).

### Model Application: Vaccine Comparison

Our model suggests that the risk of severe ADE-related disease pathology is highest when antibody titers are low or intermediate, and that the risk of ADE declines monotonically with increasing numbers of CD8+ T cells. This provides a possible explanation for the different outcomes of Dengvaxia and TAK-003 vaccine trials. Both vaccines are live attenuated flaviviruses expressing surface proteins from the four dengue strains. The vaccines differ in their internal proteins, which form the viral backbone and typically elicit CD8+ T-cell responses. The backbone used to construct the CYD-TDV vaccine is attenuated yellow fever virus (YFV-17); in contrast, TAK-003 incorporates the DENV-2 backbone. Consequently, we expect that TAK-003 generates a large dengue-specific CD8+ T-cell response [54, 32] and that CYD-TDV (Dengvaxia) generates a minimal dengue-specific T-cell response [24]. This provides a potential explanation of the higher incidence of severe disease in individuals vaccinated with CYD-TDV (Dengvaxia) as compared with unvaccinated controls [45], which was not observed for individuals vaccinated with TAK-003 [49]. The increased severity occurred about 3-4 years after vaccination of naive individuals, a timeframe that allowed waning of antibodies from a protective level to one that allowed ADE.

## 3 Materials and Methods

### Description of Model

We built a model to describe the within-host dynamics of acute dengue infections, building on earlier models of virus infection dynamics [39, 35, 7, 38] and incorporating ADE as described in Nikas et al. [33]. The model considers populations of uninfected and infected cells, virus, humoral immune responses, and CD8+ T-cell responses. Throughout the manuscript, we use “humoral immunity” to refer collectively to B cells, plasma cells, and antibodies; however, when discussing ADE and viral neutralization, we refer explicitly to antibody concentration, as this component directly mediates ADE in the model.

Viral dynamics are governed by two populations: unbound virus (*V*_0_) and virus partially bound by antibody (*V*_1_). *V*_0_ infects uninfected cells (U) at a rate proportional to viral infectivity (*β*). Because dengue virus clearance is driven primarily by immune responses rather than depletion of susceptible cells, we set *dU/dt* = 0, consistent with previously published within-host dengue models [9, 33]. *V*_1_ is more infectious than *V*_0_ and infects cells at a rate *εβ*, where *ε >* 1 represents the enhancement of infection mediated by partially bound antibody. Infected cells (I) are produced from both populations and are eliminated by CD8+ T-cell killing at rate *k*_*T*_, by innate immunity at rate *k*_*X*_, and by non-immune death at rate *d*_*I*_ . Infected cells produce new *V*_0_ at rate *p*_*V*_ . *V*_0_ is converted to *V*_1_ by antibody binding at rate *k*_0_ and is degraded at rate *d*_*V*_ . *V*_1_ is further neutralized by additional antibody binding at rate *k*_*A*_ and degraded at rate *d*_*V*_ ; fully neutralized virus is unable to infect cells and is rapidly cleared.

Immune responses are driven by virus concentration. Innate immunity (X) activates with half-saturation constant *ϕ*_*X*_ and maximum growth rate *q*. B cells expand with maximum rate *s*_*B*_ and half-saturation constant *ϕ*_*B*_; a fraction *f* differentiate into plasma cells, while the remainder die at rate *d*_*B*_. Plasma cells secrete antibodies at rate *p*_*A*_ and die at rate *d*_*P*_ ; antibodies are degraded at rate *d*_*A*_. CD8+ T cells expand with maximum rate *s*_*T*_ and half-saturation constant *ϕ*_*T*_, and are eliminated at rate *d*_*T*_ . Infection pathology is estimated as log_10_(AUC(*V*_0_ + *V*_1_)), where AUC denotes the time integral of (*V*_0_ + *V*_1_) from infection to viral clearance (resolution of infection). This approximation is consistent with clinical observations linking dengue pathology to viral load [51, 53].

The model recapitulates *in vitro* ADE dynamics in that low or intermediate antibody titers increase the number of infected cells (Fig. S1). The model is also consistent with *in vivo* NHP passive transfer experiments showing enhanced viral replication at intermediate antibody concentrations (Fig. S2) [16]. To explore the effects of prior immunity on infection outcome, we vary the initial conditions at the time of infection rather than explicitly modeling immune waning.

#### System of Differential Equations

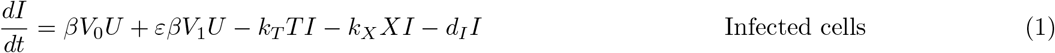

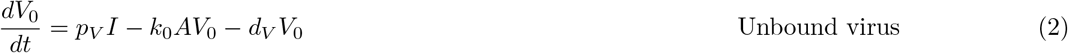

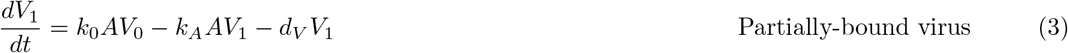

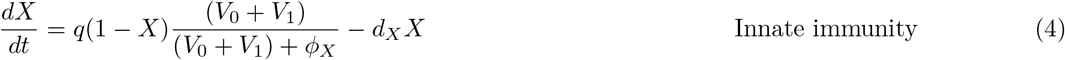

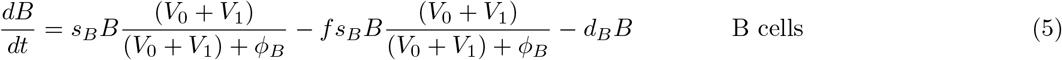

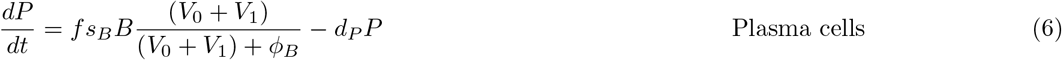

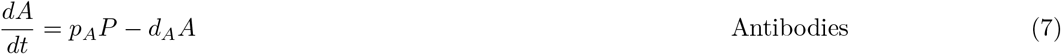

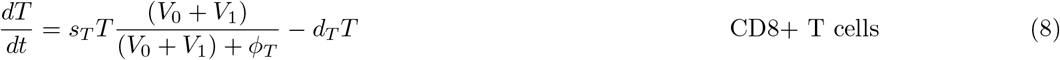

## 4 Discussion

We used our model to explore how the interplay between humoral and CD8+ T-cell immunity influences the dynamics and severity of primary and secondary dengue infections. The model suggests a potential role for CD8+ T cells in attenuating ADE under conditions in which antibody levels are low-to-intermediate (as may occur following natural infection or vaccination, when humoral immunity has waned from peak levels but remains detectable). CD8+ T-cell immunity could therefore represent an important factor that, when induced by natural infection, helps protect against ADE-enhanced severe disease as humoral immunity wanes. Because the model is intended to capture qualitative relationships between humoral and cellular immunity rather than to provide quantitative forecasts, we focus on general dynamical patterns rather than direct fits to individual patient datasets.

Vaccines, including those against dengue viruses, are typically designed with a primary focus on eliciting humoral immunity. In contrast, CD8+ T cell responses have received comparatively less attention, despite the fact that flaviviruses such as Yellow Fever Virus—and the live-attenuated YFV vaccine—elicit robust CD8+ T-cell responses [4, 13]. Recent studies have highlighted roles for CD8+ T cells in both natural dengue infection [34, 48, 14, 15] and responses to dengue vaccination [55]. Consistent with a protective role for cellular immunity, a small human challenge study demonstrated that sterilizing immunity to a related human *Orthoflavivirus* can be achieved through CD8+ T-cell responses alone, in the absence of neutralizing antibodies [28]. Together, these findings suggest that cellular immunity may be sufficient for protection against flaviviral infection in some contexts. Accordingly, we incorporated both CD8+ T-cell and humoral immune responses into our model of dengue infection.

Our results suggest that the combined effect of humoral and CD8+ immunity may help explain why severe disease is only exhibited in a subset of infections of individuals with enhancing antibody titers. In our simulations, severe disease requires not only low-to-intermediate antibody levels (necessary for ADE) but also weak CD8+ T-cell responses, which would otherwise attenuate the effects of ADE. Our models also provide a possible explanation for why the effects of ADE are more consistent and pronounced in experimental settings that do not include CD8+ T cells, such as *in vitro* cell culture experiments or *in vivo* passive antibody transfer experiments in NHPs.

The main result of our paper pertains to understanding the conditions under which vaccine-generated immunity can cause ADE. Our results provide a possible explanation for why the increased risk of severe disease following immunization with the Dengvaxia vaccine was not observed for the TAK-003 vaccine. We suggest that this may be because TAK-003 induced a stronger dengue-specific CD8+ T-cell response than CYD-TDV. More generally, our results suggest that vaccines that focus only on the generation of humoral immunity may carry a higher theoretical risk of ADE than vaccines that generate both humoral and CD8+ T-cell immunity. Our model suggests that vaccines eliciting robust CD8+ T-cell immunity with limited humoral responses may reduce the risk of ADE. For dengue and other viruses known to cause ADE (Zika [8], Ebola [47, 30], and potentially Chikungunya [31], where ADE has been proposed but is less well established), these vaccines could provide protection against severe infection without increasing the risk of severe disease. Future studies measuring both antibody titers and dengue-specific CD8+ T-cell responses prior to infection could help test the model predictions presented here.

### Caveats and Limitations

Our model is intentionally simple and does not explicitly incorporate many of the complexities that can contribute to the outcome of human dengue infections. In particular, the model is deterministic and does not include sources of host heterogeneity, such as genetic variation or aging, that are known to influence disease severity and outcomes [17, 1]. Incorporating such heterogeneity would be expected to broaden the distribution of infection outcomes but would not alter the qualitative relationships between antibody levels, CD8+ T-cell immunity, and pathology identified here.

We also do not explicitly include how disease severity may be affected by the infecting serotype or strain, or by interactions between viral genotype and pre-existing immunity [46, 36, 40, 9, 11, 45, 52, 50, 2]. While these factors could influence the trajectory of immune boosting and waning (Fig. 5), we expect the qualitative results to remain unchanged. A more nuanced framework would be required to provide a quantitatively accurate description of how specific serotype and strain combinations affect pathology. Finally, although our hypothesis is consistent with existing vaccine trial results, CD8+ T-cell responses may, in some contexts, contribute to immunopathology. Further studies are needed to determine the conditions under which CD8+ T-cell responses are protective or pathogenic during dengue infection, and whether additional factors contributed to severe disease outcomes observed in some CYD-TDV recipients.

### Future Directions

Our current model is intentionally simple, and as additional data become available it could be refined. Future models could consider specific effects of serotypes and the number of prior infections, as well as interactions between strains. Future population-level modeling could also account for boosting, waning, immunological heterogeneity between individuals, and the age structure of the population. Understanding the population-level impacts of antibody-dependent enhancement may require modeling B-cell and T-cell waning separately. These waning dynamics have previously been described in the context of other pathogens [42, 5]. On a practical level, our results suggest that vaccines, particularly dengue vaccines, should consider the potential benefit of eliciting CD8+ T-cell responses.

## Acknowledgments

We thank colleagues for helpful discussions and feedback on the modeling framework and interpretation of the results. We acknowledge the following support: U01 AI150747 (RA). The funders had no role in study design, data collection and interpretation, or the decision to submit the work for publication.

## Notes

### Competing Interest Statement

The authors have declared no competing interest.

## References

[1] Abel, L. / Casanova, J.-L. (2024). The microbe, the infection enigma, and the host. Annual Review of Microbiology 78: 103–124. 10.1146/annurev-micro-092123-022855

[2] Aggarwal, C., Ahmed, H., Sharma, P., et al. (2024). Severe disease during both primary and secondary dengue virus infections in pediatric populations. Nature Medicine 30(3): 670–674. 10.1038/s41591-024-02798-x

[3] Aguiar, M., Stollenwerk, N., Halstead, S.B. (2016). The risks behind Dengvaxia recommendation. Lancet Infectious Diseases 16(8): 882–883.10.1016/S1473-3099(16)30168-2

[4] Akondy, R.S., Monson, N.D., Miller, J.D., Edupuganti, S., Teuwen, D., Wu, H., Quyyumi, F., Garg, S., Altman, J.D., Del Rio, C., Keyserling, H.L., Ploss, A., Rice, C.M., Orenstein, W.A., Mulligan, M.J., Ahmed, R. (2009). The yellow fever virus vaccine induces a broad and polyfunctional human memory CD8+ T cell response. Journal of Immunology 183(12): 7919–7930. 10.4049/jimmunol.0803903

[5] Amanna, I.J., Carlson, N.E., Slifka, M.K. (2007). Duration of humoral immunity to common viral and vaccine antigens. NEJM 357(19): 1903–1915. 10.1056/NEJMoa066092

[6] Antia, R., Levin, B.R., May, R.M. (1994). Within-host population dynamics and the evolution and maintenance of microparasite virulence. The American Naturalist 144(3): 457–472. 10.1086/285686

[7] Baccam, P., Beauchemin, C., Macken, C.A., Hayden, F.G., Perelson, A.S. (2006). Kinetics of influenza A virus infection in humans. Journal of Virology 80(15): 7590–7599. 10.1128/JVI.01623-05

[8] Bardina, S.V., Bunduc, P., Tripathi, S., Duehr, J., Frere, J.J., Brown, J.A., Nachbagauer, R., Foster, G.A., Krysztof, D., Tortorella, D., Stramer, S.L., García-Sastre, A., Krammer, F., Lim, J.K. (2017). Enhancement of Zika virus pathogenesis by preexisting antiflavivirus immunity. Science 356(6334): 175–180. 10.1126/science.aal4365

[9] Ben-Shachar, R., Schmidler, S., & Koelle, K. (2016). Drivers of inter-individual variation in dengue viral load dynamics. PLOS Computational Biology 12(11): e1005194. 10.1371/journal.pcbi.1005194

[10] Butrapet, S., Huang, C.Y., Pierro, D.J., Bhamarapravati, N., Gubler, D.J., & Kinney, R.M. (2000). Attenuation markers of a candidate dengue type 2 vaccine virus, strain 16681 (PDK-53), are defined by mutations in the 5 noncoding region and nonstructural proteins 1 and 3. Journal of Virology 74(7): 3011–3019. 10.1128/JVI.74.7.3011-3019.2000

[11] Clapham, H.E., Cummings, D.A.T., & Johansson, M.A. (2017). Immune status alters the probability of apparent illness due to dengue virus infection: Evidence from a pooled analysis across multiple cohort and cluster studies. PLOS Neglected Tropical Diseases 11(9): e0005926. 10.1371/journal.pntd.0005926

[12] Dejnirattisai, W., Jumnainsong, A., Onsirisakul, N., Fitton, P., Vasanawathana, S., Limpitikul, W., et al. (2010). Cross-reacting antibodies enhance dengue virus infection in humans. Science 328: 745–748. 10.1126/science.1185181

[13] Fuertes Marraco, S.A., Soneson, C., Cagnon, L., Gannon, P.O., Allard, M., Abed Maillard, S., Montandon, N., Rufer, N., Waldvogel, S., Delorenzi, M., & Speiser, D.E. (2015). Long-lasting stem cell-like memory CD8+ T-cells with a naive-like profile upon yellow fever vaccination. Science Translational Medicine 7(282): 282ra48. 10.1126/scitranslmed.aaa3700

[14] Friberg, H., Martinez, L.J., Lin, L., Blaylock, J.M., De La Barrera, R.A., Rothman, A.L., et al. (2020). Cell-mediated immunity from purified inactivated DENV-1 vaccine. mSphere 5(1): e00671–19. 10.1128/mSphere.00671-19

[15] Gálvez, R.I., Martínez-Pérez, A., Escarrega, E.A., Singh, T., Zambrana, J.V., Balmaseda, Á., Harris, E., Weiskopf, D. (2025). Frequency of dengue virus-specific T-cells relates to infection outcomes. JCI Insight 10(4): e179771. 10.1172/jci.insight.179771

[16] Goncalvez, A.P., Engle, R.E., St. Claire, M., Purcell, R.H., Lai, C. (2007). Monoclonal antibody-mediated enhancement of dengue infection. PNAS 104(22): 9422–9427. 10.1073/pnas.0703498104

[17] Glynn, J.R. & Moss, P.A.H. (2020). Systematic analysis of infectious disease outcomes by age shows lowest severity in school-age children. Scientific Data 7(1): 329. 10.1038/s41597-020-00668-y

[18] Haider, N., Hasan, M.N., Onyango, J., Billah, M., Khan, S., Papakonstantinou, D., et al. (2025). Global dengue epidemic with record 14M cases in 2024. International Journal of Infectious Diseases 158: 107940. 10.1016/j.ijid.2025.107940

[19] Hadinegoro, S.R., Arredondo-García, J.L., Capeding, M.R., Deseda, C., Chotpitayasunondh, T., Dietze, R., et al. (2015). Efficacy & long-term safety of dengue vaccine in endemic regions. NEJM 373(13): 1195–1206. 10.1056/NEJMoa1506223

[20] Halstead, S.B., Nimmannitya, S., Cohen, S.N. (1967). Pathogenesis of dengue hemorrhagic fever: immune response & virus recovered. Yale J Biol Med 42: 311–328.

[21] Halstead, S.B., Shotwell, H., Casals, J. (1973a). Pathogenesis in monkeys: primary infection. J Infect Dis 128: 7–14. 10.1093/infdis/128.1.7

[22] Halstead, S.B., Shotwell, H., Casals, J. (1973b). Pathogenesis in monkeys: heterologous infection. J Infect Dis 128: 15–22. 10.1093/infdis/128.1.15

[23] Halstead, S.B., O’Rourke, E.J. (1977). Antibody-enhanced dengue virus infection. J Exp Med 146: 201–217. 10.1084/jem.146.1.201

[24] Harenberg, A., Begue, S., Mamessier, A., Gimenez-Fourage, S., Ching Seah, C., Wei Liang, A., Li, J., Nien-Yung Toh, X., Nosten, F., Rivino, L., Tan, S.B., Bertoletti, A., Cooke, A., Sutherland, C.J.R., Carre, H., Bouckenooghe, A., Wartel, T.A., Guy, B., Lang, J. (2013). Persistence of Th1/Tc1 responses one year after tetravalent dengue vaccination in adults and adolescents in Singapore. Human Vaccines & Immunotherapeutics 9(11): 2317–2330. 10.4161/hv.25562

[25] Hawkes, R.A. (1964). Enhancement of arbovirus infectivity by antibodies. Aust J Exp Biol Med Sci 42: 465–482. 10.1038/icb.1964.44

[26] Hawkes, R.A., Lafferty, K.J. (1967). Enhancement of virus infectivity by antibody. Virology 33: 250–261. 10.1016/0042-6822(67)90144-4

[27] Hou, J., Ye, W., Chen, J. (2022). Development & challenges of tetravalent live-attenuated dengue vaccines. Front Immunol 13: 840104. 10.3389/fimmu.2022.840104

[28] Kalimuddin, S., Tham, C.Y.L., Chan, Y.F.Z., et al. (2025). Vaccine-induced T-cells control Orthoflavivirus infection without nAbs. Nature Microbiology 10: 374–387. 10.1038/s41564-024-01903-7

[29] Katzelnick, L.C., Gresh, L., Halloran, M.E., Mercado, J.C., Kuan, G., Gordon, A., et al. (2017). Antibody-dependent enhancement of severe dengue disease. Science 358: 929–932. 10.1126/science.aan6836

[30] Kuzmina, N.A., Younan, P., Gilchuk, P., Santos, R.I., Flyak, A.I., Ilinykh, P.A., et al. (2018). ADE of Ebola virus by survivor antibodies. Cell Reports 24: 1802–1815.e5. 10.1016/j.celrep.2018.07.035

[31] Lum, F.M., Couderc, T., Chia, B.S., Ong, R.Y., Her, Z., Chow, A., et al. (2018). ADE worsens chikungunya severity. Scientific Reports 8: 1860. 10.1038/s41598-018-20305-4

[32] Mandarić, S., Friberg, H., Sáez-Llorens, X., Borja-Tabora, C., Biswal, S., Escudero, I., et al. (2024). Long-term T cell response & safety of tetravalent dengue vaccine. NPJ Vaccines 9: 192. 10.1038/s41541-024-00967-0

[33] Nikas, A., Ahmed, H., Moore, M.R., Zarnitsyna, V.I., Antia, R. (2023). When does humoral memory enhance infection? PLoS Comp Biol 19(8): e1011377. 10.1371/journal.pcbi.1011377

[34] Elong Ngono, A., Chen, H.W., Tang, W.W., Joo, Y., King, K., Weiskopf, D., Sidney, J., Sette, A., Shresta, S. (2016). Protective role of cross-reactive CD8 T-cells against dengue virus infection. EBioMedicine 13: 284–293. 10.1016/j.ebiom.2016.10.006

[35] Nowak, M.A., Perelson, A.S. (1994). HIV replication and immune control dynamics. Seminars in Virology 5: 165–180. 10.1006/smvy.1994.1014

[36] OhAinle, M., Balmaseda, A., Macalalad, A.R., Tellez, Y., Zody, M.C., Saborío, S., et al. (2011). Dengue severity driven by viral genetics × immunity interactions. Sci Transl Med 3(114): 114ra128. 10.1126/scitranslmed.3003084

[37] Patel, S.S., Rauscher, M., Kudela, M., Pang, H. (2023). Clinical safety experience of TAK-003 for dengue fever: A new tetravalent live attenuated vaccine candidate. Clinical Infectious Diseases 76(3): e1350–e1359. 10.1093/cid/ciac418

[38] Pawelek, K.A., Huynh, G.T., Quinlivan, M., Cullinane, A., Rong, L., Perelson, A.S. (2012). Influenza within-host dynamics with immune response. PLoS Comp Biol 8(6): e1002588. 10.1371/journal.pcbi.1002588

[39] Perelson, A.S., Wiegel, F.W. (1992). Design principles for immune recognition. Ann Rev Biophys Biomol Struct 21: 331–360. 10.1146/annurev.bb.21.060192.001555

[40] Reich, N.G., Shrestha, S., King, A.A., Rohani, P., Lessler, J., Kalayanarooj, S., Yoon, I.-K., Gibbons, R.V., Burke, D.S., & Cummings, D.A.T. (2013). Interactions between serotypes of dengue highlight epidemiological impact of cross-immunity. Journal of The Royal Society Interface 10(86): 20130414. 10.1098/rsif.2013.0414

[41] Rivera, L., Biswal, S., Sáez-Llorens, X., Reynales, H., López-Medina, E., Borja-Tabora, C., et al. (2022). Three-year efficacy & safety of TAK-003. Clin Infect Dis 75(1): 107–117. 10.1093/cid/ciab864

[42] Saha, A., Ahmed, H., Hirst, C., Koelle, K., Handel, A., Teunis, P., Antia, R. (2025). Quantifying the waning of humoral immunity. Immunity 58(12): 3144–3152.e2. 10.1016/j.immuni.2025.11.007

[43] Salje, H., Cummings, D.A.T., Rodriguez-Barraquer, I., Katzelnick, L.C., Lessler, J., Klungthong, C., et al. (2018). Antibody dynamics & dengue risk. Nature 557: 719–723. 10.1038/s41586-018-0157-4

[44] Salje, H., Alera, M.T., Chua, M.N., Hunsawong, T., Ellison, D., Srikiatkhachorn, A., Jarman, R.G., Gromowski, G.D., Rodriguez-Barraquer, I., Cauchemez, S., Cummings, D.A.T., Macareo, L., Yoon, I.K., Fernandez, S., Rothman, A.L. (2021). Evaluation of the extended efficacy of the Dengvaxia vaccine against symptomatic and subclinical dengue infection. Nature Medicine 27(8): 1395–1400. 10.1038/s41591-021-01392-9

[45] Sridhar, S., Luedtke, A., Langevin, E., Zhu, M., Bonaparte, M., Machabert, T., Savarino, S., Zambrano, B., Moureau, A., Khromava, A., Moodie, Z., Westling, T., Mascareñas, C., Frago, C., Cortés, M., Chansinghakul, D., Noriega, F., Bouckenooghe, A., Chen, J., Ng, S.P., Gilbert, P.B., Gurunathan, S., DiazGranados, C.A. (2018). Effect of dengue serostatus on dengue vaccine safety and efficacy. New England Journal of Medicine 379(4): 327–340. 10.1056/NEJMoa1800820

[46] Sierra, B., Perez, A.B., Vogt, K., Garcia, G., Schmolke, K., Aguirre, E., et al. (2010). Immune regulation vs inflammation in secondary heterologous dengue. Cell Immunol 262: 134–140. 10.1016/j.cellimm.2010.02.005

[47] Takada, A., Feldmann, H., Ksiazek, T.G., Kawaoka, Y. (2003). ADE of Ebola virus infection. J Virol 77(13): 7539–7544. 10.1128/JVI.77.13.7539-7544.2003

[48] Tian, Y., Grifoni, A., Sette, A., Weiskopf, D. (2019). Human T cell response to dengue. Front Immunol 10: 2125. 10.3389/fimmu.2019.02125

[49] Tricou, V., Yu, D., Reynales, H., Biswal, S., Sáez-Llorens, X., Sirivichayakul, C., et al. (2024). 4.5-year TAK-003 efficacy & safety. Lancet Global Health 12: e257–e270. 10.1016/S2214-109X(23)00522-3

[50] Tsang, T.K., Ghebremariam, S.L., Gresh, L., et al. (2019). Effects of infection history on dengue virus infection and pathogenicity. Nature Communications 10: 1246. 10.1038/s41467-019-09193-y

[51] Vaughn, D.W., Green, S., Kalayanarooj, S., Innis, B.L., Nimmannitya, S., Suntayakorn, S., Endy, T.P., Raengsakulrach, B., Rothman, A.L., Ennis, F.A., Nisalak, A. (2000). Dengue viremia titer, antibody response pattern, and virus serotype correlate with disease severity. Journal of Infectious Diseases 181(1): 2–9. 10.1086/315215

[52] Vicente, C.R., Herbinger, K.H., Fröschl, G., et al. (2016). Serotype influences on dengue severity in Brazil. BMC Infect Dis 16: 320. 10.1186/s12879-016-1668-y

[53] Waggoner, J.J., Katzelnick, L.C., Burger-Calderon, R., Gallini, J., Moore, R.H., Kuan, G., Balmaseda, A., Pinsky, B.A., Harris, E. (2020). Antibody-dependent enhancement of severe disease is mediated by serum viral load in pediatric dengue virus infections. Journal of Infectious Diseases 221(11): 1846–1854. 10.1093/infdis/jiz618

[54] Waickman, A.T., Victor, K., Li, T., Hatch, K., Rutvisuttinunt, W., Medin, C., et al. (2019). Heterogeneity of DENV vaccine-elicited T-cells via scRNAseq. Nature Communications 10: 3666. 10.1038/s41467-019-11634-7

[55] Waickman, A.T., Friberg, H., Gromowski, G.D., Rutvisuttinunt, W., Li, T., Siegfried, H., et al. (2021). Temporal scRNAseq of PBMC in experimental & natural DENV-1 infection. PLOS Pathogens 17: e1009240. 10.1371/journal.ppat.1009240

[56] Wilder-Smith, A., et al. (2019). Deliberations of the Strategic Advisory Group of Experts on Immunization on the use of CYD-TDV dengue vaccine. Lancet Infectious Diseases 19: e31–e38. 10.1016/S1473-3099(18)30494-8

