## Supplemental Material for "CD8+ T cells and Humoral Immunity Influence the Development of Antibody-Dependent Enhancement: Implications for Vaccine Design"

Drake et al., 2026

#### Supplemental Table S1. Model summary: state variables, parameters, and initial conditions

| **State Variables and Their Interpretations** | | |
|---|---|---|
| **State Variable** | **Interpretation** | **Units** |
| $I$ | Proportion of target-cells infected | – |
| $U$ | Proportion of target-cells uninfected | – |
| $V_0$ | Virus not bound by antibodies, infectivity $\beta$ | VU |
| $V_1$ | Virus partially bound by antibodies, infectivity $\beta * \varepsilon$ | VU |
| $T$ | CD8+ T-cell immunity | CU |
| $X$ | Innate immune response | RU |
| $B$ | Pathogen-specific B-cells | CU |
| $P$ | Antibody-secreting plasma cells | CU |
| $A$ | Antibody concentration | AU |
| **Model Parameters** | | |
| **Parameter** | **Interpretation** | **Default Value** |
| $\beta$ | Infectivity of unbound virus | 5 CU/VU/day |
| $\varepsilon$ | Increased infectivity of semi-bound virus | 1.5 |
| $k_T$ | Killing rate of infected cells by CD8+ T cells | 0.002 CU/day |
| $k_X$ | Killing rate by innate immunity | 4 CU/RU/day |
| $d_I$ | Death rate of infected cells | 1/day |
| $p_V$ | Virus production rate | 20 VU/CU/day |
| $k_0$ | Antibody binding rate ($V_0 \to V_1$) | 0.5/AU/day |
| $d_V$ | Virus clearance rate | 15/day |
| $k_A$ | Neutralization rate of semi-bound virus | 0.01/AU/day |
| $s_T$ | CD8+ T-cell proliferation rate | 1/day |
| $d_T$ | CD8+ T-cell death rate | 0/day |

#### Supplemental Figures

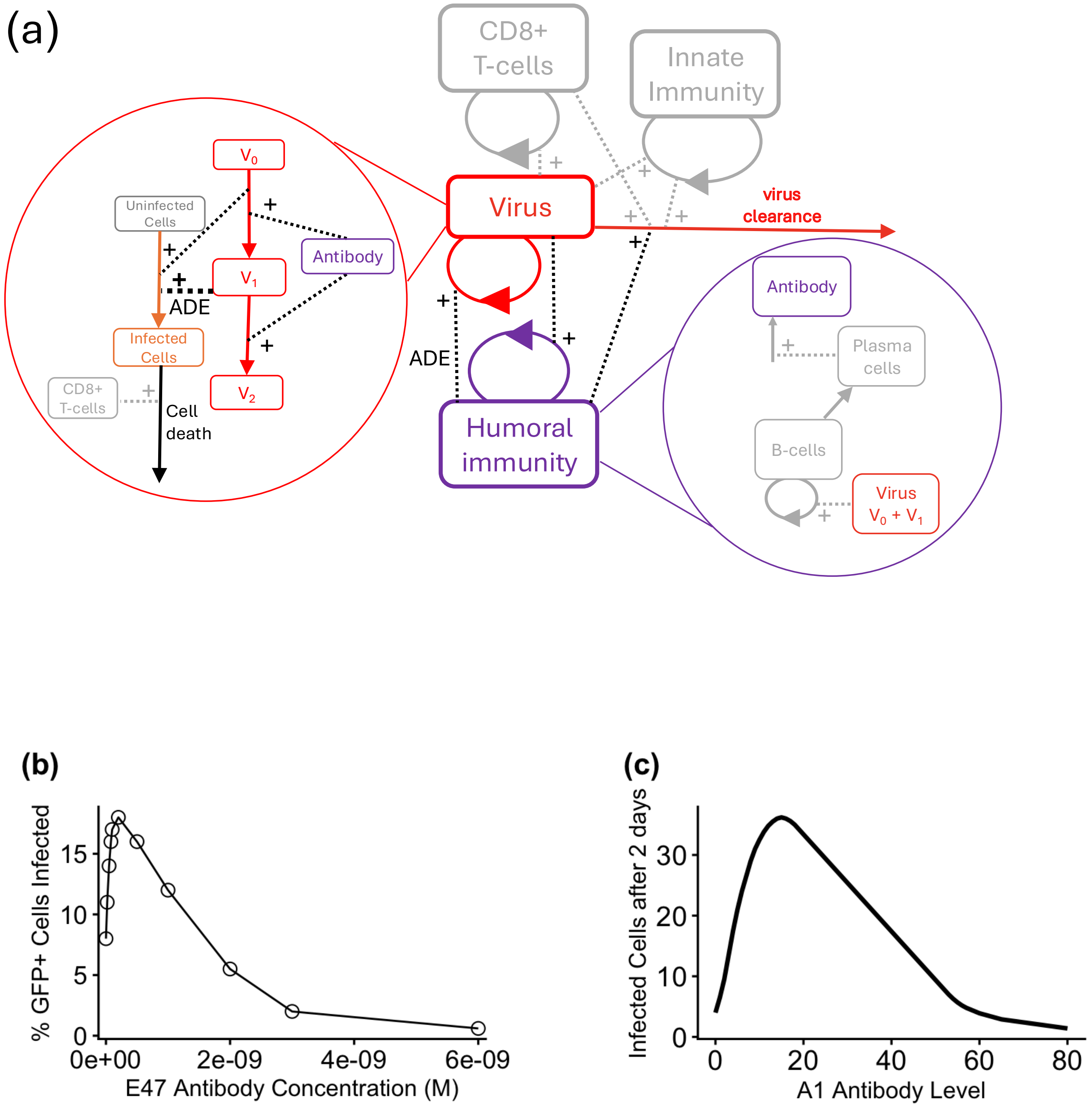


Fig. S1: (a) Components of the model (shown in color) used to test whether it captures the *in vitro* dynamics of antibody-dependent enhancement. (b) Experimental results demonstrating *in vitro* ADE in the absence of immunity. Data digitized from panel 4A of [1]. (c) Model results under the same conditions.

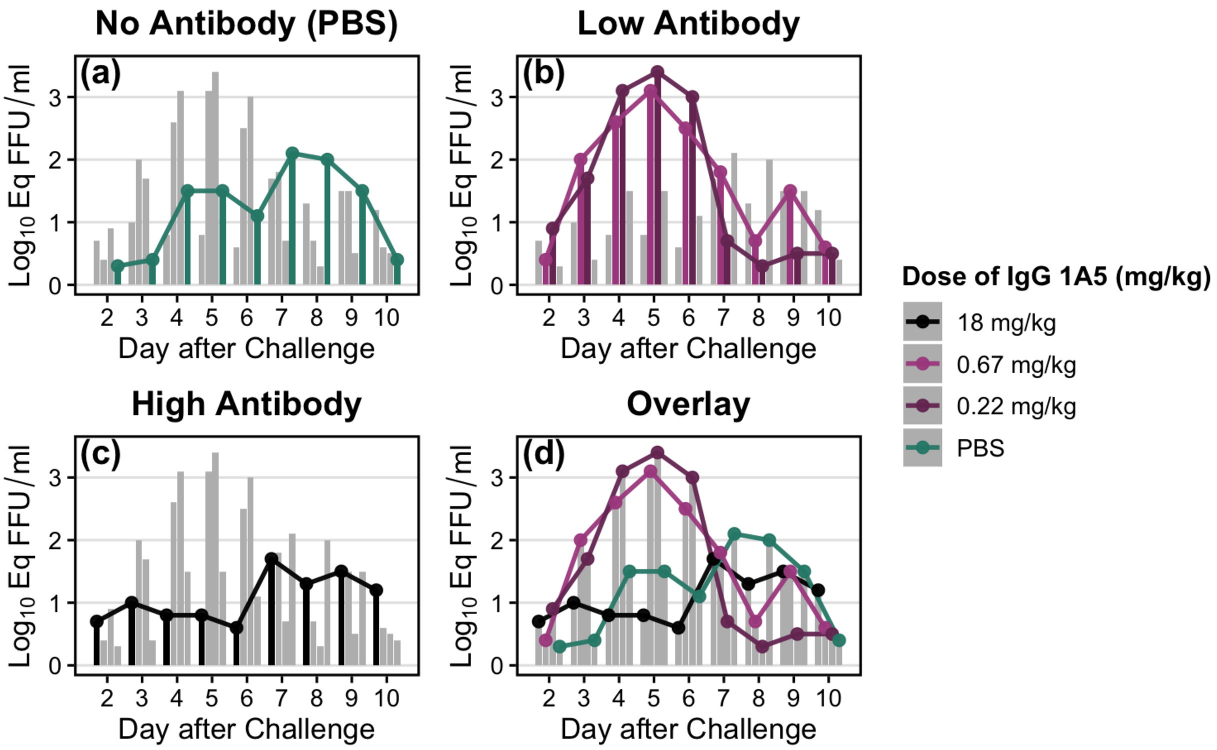


Fig. S2: **Dengue Infection Course in NHPs After IgG 1A5 Pre-Dose** Our within-host model captures *in vivo* dynamics of antibody-dependent enhancement in non-human primates infected with dengue just after passive antibody transfer of dengue-specific antibodies. This figure shows the viral dynamics of dengue infections in non-human primates (NHPs) that were administered different doses of anti-dengue antibodies just before infection (a) shows the mean viral titers across the course of infection for animals that did not receive any passive antibody before infection (n = 3) (b) shows mean viral titers for animals that received a low (0.22 mg/kg) or intermediate (0.67 mg/kg) dose of anti-dengue antibody just before dengue infection (n = 3 for each dose group) (c) shows mean viral titers for animals that received a high dose of anti-dengue antibody (18 mg/kg) just before dengue infection (n = 3). In panel (d) antibody dosage groups are overlaid, demonstrating that, compared with animals that received no, low, or intermediate amounts of passively transferred antibodies, viral replication is substantially suppressed in animals that received the high antibody dose. NHPs that received low- or intermediate-antibody doses had higher viral replication than those that received no passively transferred antibody.

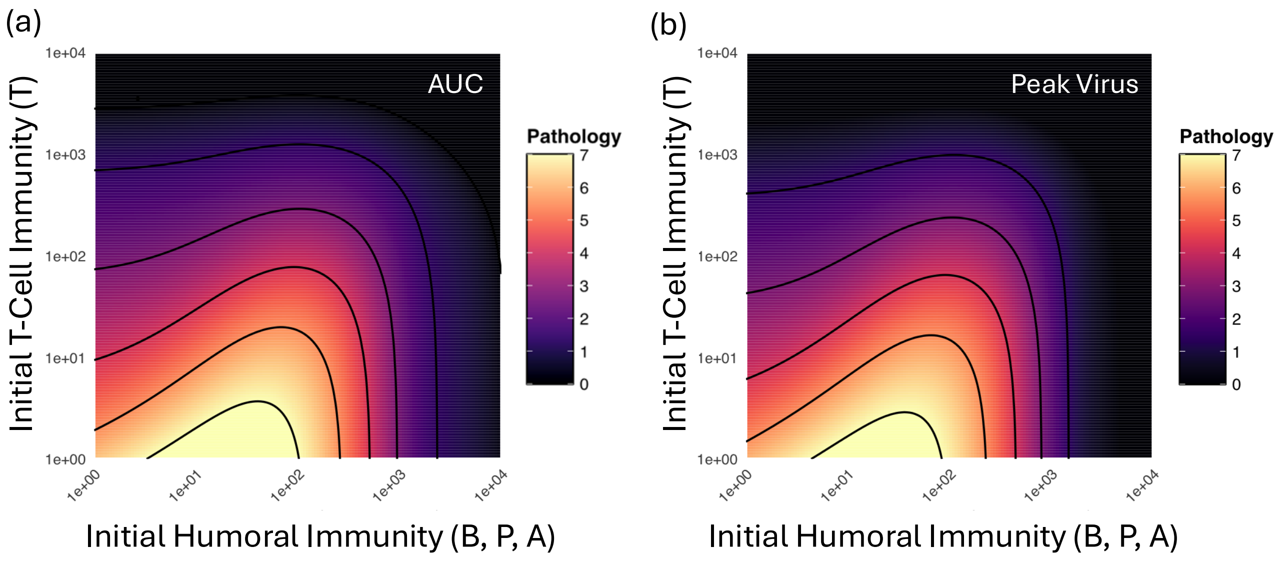


Fig. S3: **The effect of pre-existing immunity on two metrics of pathology** In Panel A, pathology is approximated by $\log_{10}$ of the area under the curve (AUC) of the infectious virus ($V0+V_1$) during the infection course. Panel B shows simulation results for a potential alternative metric of pathology, Peak Viral Load. In our model, infections almost always resolve within 30 days; both metrics shown are calculated using a 30-day time horizon (most infections resolve much more quickly, within 12–17 days). The infected person starts with different levels of B-cell immunity (on the x-axis) and T-cell immunity (on the y-axis). The color at each position on the heatmap indicates the pathological severity of the infection, with the starting conditions indicated by the x-axis (B-cell immunity) and y-axis (T-cell immunity)—darker colors indicate less severe infections, and lighter colors indicate more severe infections. Both pathology metrics show that infections are enhanced by prior immunity when individuals begin infections with low/intermediate B-cell immunity and no CD8+ T-cell immunity, and that the presence of specific CD8+ T cells before infection protects against antibody-dependent enhancement in a dose-dependent manner.

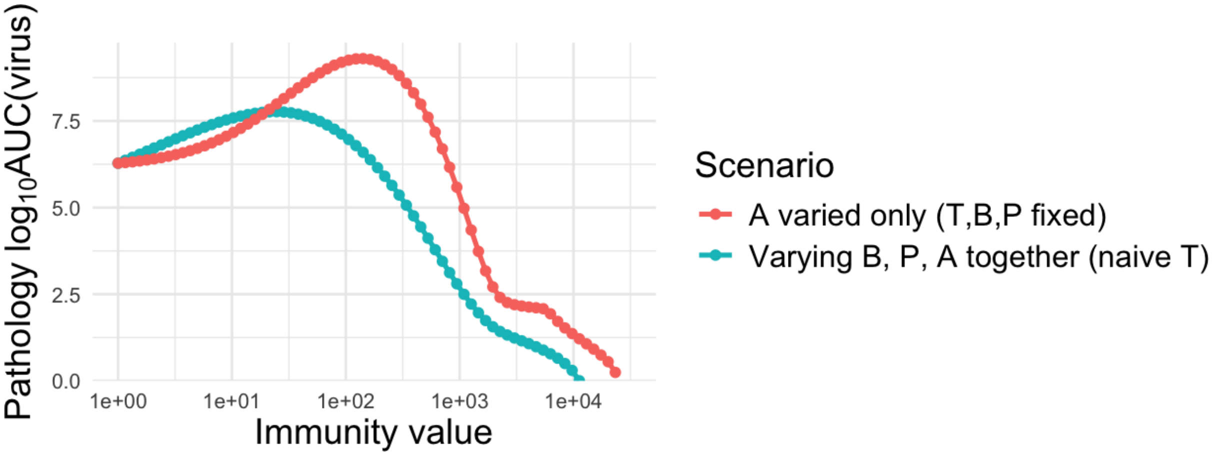


Fig. S4: **Humoral immunity and antibody versus infection pathology** Infection pathology (y-axis) is shown as a function of the pre-infection levels of humoral immunity (blue line) or passively transferred antibody (red line). The red curve shows the effect on infection pathology of increasing antibody alone; higher antibody values increase infection pathology to a greater magnitude and over a broader range than are reached when increasing humoral immunity. Increasing antibody alone produces a marked increase in pathology at low-to-intermediate antibody levels, consistent with substantial enhancement of infection pathology resulting from passive antibody transfer. These results are consistent with the observed pattern that antibody-enhanced severe disease is a rare outcome in cases of natural infection and immunity—specifically that increasing humoral immunity (B, P, and A) causes an increase in infection pathology over a much smaller range and at a much lower magnitude than increasing just the amount of passive antibody transferred (A).

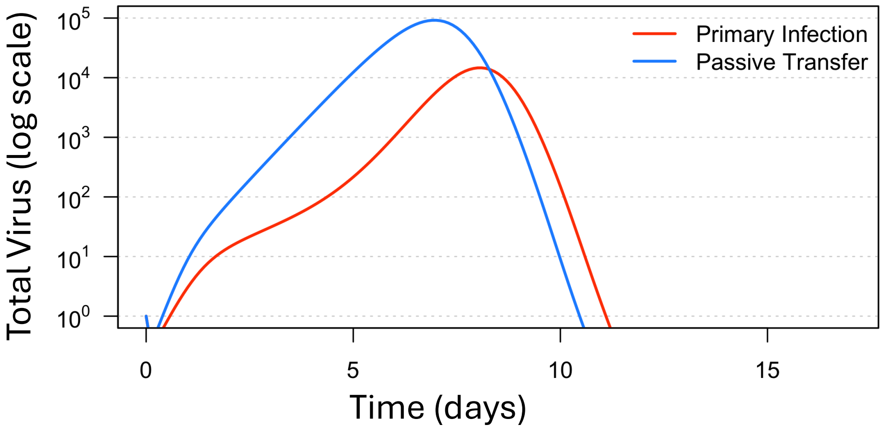


Fig. S5: **Viral dynamics in primary infection vs. passive antibody transfer** Passive antibody transfer increases the maximum viral titer observed during infection by about 10-fold compared to primary infection without passively transferred antibody.

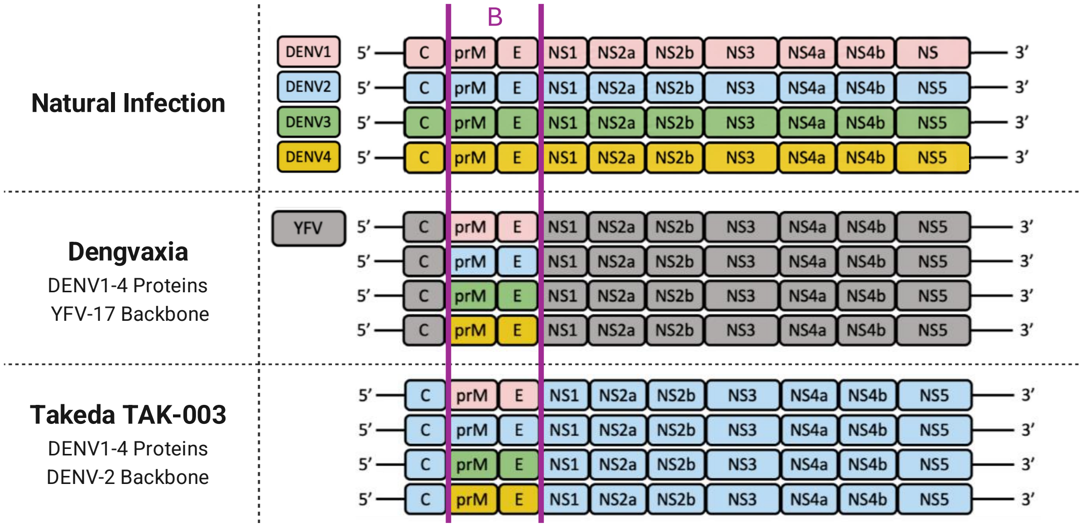


Fig. S6: Schematic comparing epitopes present in natural infection to those in each of the two licensed dengue vaccines, CYD-TDV (Dengvaxia) and TAK-003 (Qdenga). CYD-TDV was made using dengue-specific PrM and E proteins and a yellow fever backbone. TAK-003 also incorporated dengue-specific PrM and E proteins, but was on a DENV-2 backbone. While both vaccines elicited a robust humoral response, only TAK-003 generated a long-lived, dengue-specific CD8+ T-cell response [2]. In natural dengue infection, CD8+ T cells preferentially target NS3 and NS5 epitopes, while B-cell and $CD4^+$ T-cell epitopes are targeted towards recognition of envelope, capsid, and NS1 proteins [3].
*Attribution:* Adapted from Hou, Ye, and Chen (2022), *Frontiers in Immunology* 13:840104 (`https://www.frontiersin.org/journals/immunology/articles/10.3389/fimmu.2022.840104/full`), licensed under CC BY 4.0 (`https://creativecommons.org/licenses/by/4.0/`). Changes: additional labels and lines added.

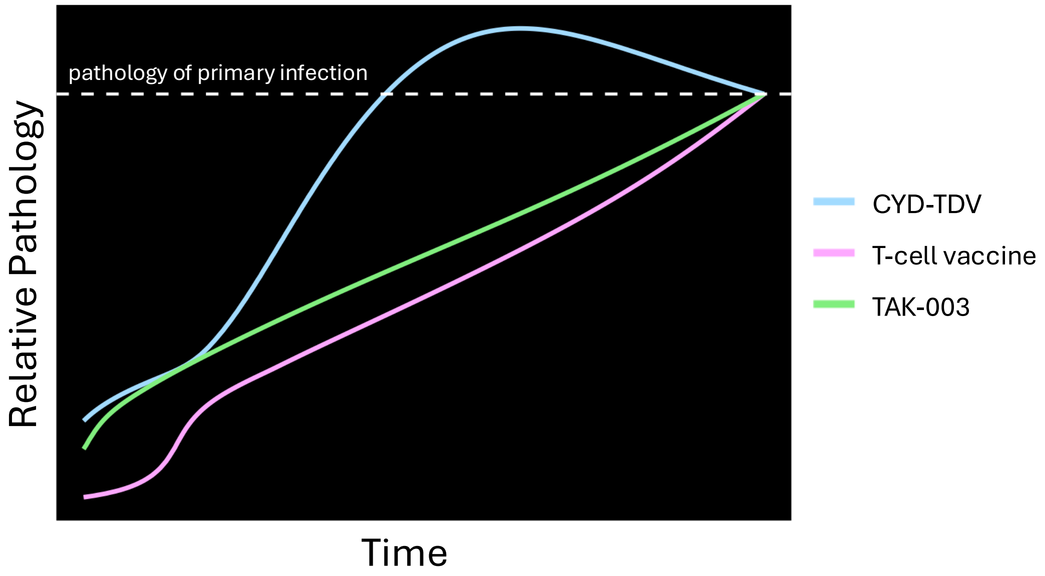


Fig. S7: **Pathology as vaccine immunity wanes** Using the waning trajectories of vaccine-generated immunity for individuals vaccinated with CYD-TDV (Dengvaxia), TAK-003 (Qdenga), or a T-cell-focused vaccine, as shown in Fig. 6, we simulate the pathology of a secondary infection at different times during the waning of vaccine-generated immunity. The dashed, white horizontal line indicates the level of pathology predicted for a primary infection. All vaccination strategies initially lower pathology and will eventually wane to a naive state. Both T-cell vaccination and TAK-003 reduce pathology compared with primary infection. Consistent with the CYD-TDV (Dengvaxia) trial results, the pathology of infection among Dengvaxia vaccine recipients is higher than that of primary infection in the 4th to 5th year after vaccination.

### References

[1] Dejnirattisai, W., Jumnainsong, A., Onsirisakul, N., Fitton, P., Vasanawathana, S., Limpitikul, W., et al. (2010). Cross-reacting antibodies enhance dengue virus infection in humans. *Science* 328: 745–748. `https://doi.org/10.1126/science.1185181`

[2] Hou, J., Ye, W., Chen, J. (2022). Development & challenges of tetravalent live-attenuated dengue vaccines. *Front Immunol* 13: 840104. `https://doi.org/10.3389/fimmu.2022.840104`

[3] Rivino, L., Kumaran, E.A., Jovanovic, V., Nadua, K., Teo, E.W., Pang, S.W., Teo, G.H., Gan, V.C., Lye, D.C., Leo, Y.S., Hanson, B.J., Smith, K.G., Bertoletti, A., Kemeny, D.M., MacAry, P.A. (2013). Differential targeting of viral components by CD4+ versus CD8+ T lymphocytes in dengue virus infection. *Journal of Virology* 87(5): 2693–2706. `https://doi.org/10.1128/JVI.02675-12`
